# Effects of Sodium tanshinone IIA sulfonate on H9C2 cardiomyocyte injury induced by sodium arsenite

**DOI:** 10.1101/2020.02.14.948505

**Authors:** Heli Xu, Xinsheng Zhang, Mengyao Liu, Jiayu Li, Wenjing Dou, Xiyan Wang, Hongyou Tan, Jian Ding, Juhua Xie, Yue Zhang

## Abstract

Arsenic is a grade I human carcinogen that can cause kinds of damage to the body. The heart is one of the main target organs of arsenic damage, but the preventive and curative measures underlying arsenic poisoning are not clear. To investigate the effect of tanshinone IIA sulfonate (STS) on the injury of H9C2 cardiomyocytes induced by NaAsO2,H9C2 cardiomyocyteswere divided into four groups where control group with general medium, arsenic (As) group received sodium arsenite (NaAsO2)-containing general medium, STS pretreatment+As group and finally the STS alone group. Compared with the control group, different concentrations of STS had no significant effect on the cell viability (n = 6, P > 0.05); Compared with the control group, there was no significant change in the viability of cells treated with arsenic for 12 h after STS pretreatment for 6 h (n = 6, P > 0.05),the viability of cells treated with NaAsO2 alone was decreased(n = 6, P < 0.05);Compared with the control group, the cell viability was decreased after STS pretreatment for 12h and arsenic for 24h, the cell viability of NaAsO2 group was also decreased, the viability of NaAsO2 cells was lower than that of STS pretreated for 12h with NaAsO2 for 24h(n=6,p<0.05).The content of Caspase 3/7 in the NaAsO2 group was higher than that in the STS and NaAsO2 group and the Control group and STS group after 12h, the differences were statistically significant(n=6,p<0.05).After STS pretreatment for 12h and NaAsO2 for 24 h, there was no significant difference in Caspase 8 content between NaAsO2 group and other groups(n=6,P>0.05) As a result, Sodium tanshinone IIA sulfonate can attenuate the injury of H9C2 cardiomyocytes induced by sodium arsenite.

## Introduction

TANIIA is one of the components extracted from Salvia Miltiorrhiza Bunge, but it is not easily absorbed through intestinal tract.And for that, Sodium tanshinone IIA sulfonate (STS) was invented to improve its bioavailability.STS, which induces Vasodilation, inhibits inflammation, prevents arteriosclerosis, heart damage and hypertrophy, is considered to be a promising natural heart protective drug.^1,2)^Arsenic is a highly toxic and widely distributed metallic element that has been used to treat tumors for more than 2,000 years, Especially for Acute promyelocytic leukemia.However, long-term exposure to arsenic can lead to various diseases, especially to cardiovascular system. Arsenic can cause many kinds of cardiac damage, including electrophysiological changes and myocardial necrosis.^3,4)^At present, there is not a good drug to antagonize the cardiac toxicity of arsenic, so this study is to investigate the effect of Sodium tanshinone IIA sulfonate on the injury of rat H9C2 cardiomyocytes induced by NaAsO2, it provides a new way to prevent and treat the cardiac toxicity of arsenic.

## Materials and methods

### Cells, Reagents

H9C2 rat cardiomyocyte strain(Shanghai Fumeng Gene Biotechnology company); DMEM Culture Medium(Hyclone company); FBS(Hyclone Corporation); Trypsin (Lifescience company); DPBS (Hyclone company); sodium arsenite (Analytical reagent, Fluka company); Tanshinone IIA sodium sulfonate(Shanghai Shifeng Biotechnology company); CCK-8 Kit (Dojindo company); caspase8 Kit (Promega company); Caspase3 / 7 Kit (Promega company).

### Cell culture

Rat H9C2 cardiomyocytes were cultured in DMEM containing 10% fetal bovine serum (FBS) at 37 °C with a volume fraction of 5% CO2. The cells were digested with 0.25% trypsin and terminated with DMEM containing 10% FBS. The passage ratio was 1:3.

### CCK-8 was used to detect the viability of cells treated with STS alone

The cells were inoculated into 96-well plates with 2000 cells / well, 6 samples in each group, then cultured in DMEM without serum for 12h. The cells were treated with 0 / 10 / 20 / 30 / 40 / 50 / 100 uMol / L STS for 12h, then the cell viability was measured by CCK-8 Kit.

### CCK-8 was used to detect the viability of cells pretreated with STS for 6h and treated with NaAsO2 for 12h

The cells were divided into control group, treated with NaAsO2 alone, the concentrations were 15umol/L、STS was pretreated for 6h and NaAsO2 was added for 12h, the concentrations of STS were 10 / 20 / 30 / 40 / 50umol/L,then the cell viability was measured by CCK-8 Kit.

### CCK-8 was used to detect the viability of cells pretreated with STS for 12h and treated with NaAsO2 for 24h

The cells were divided into control group, treated with NaAsO2 alone and the concentrations were 10umol/L、STS was pretreated for 12h and NaAsO2 was added for 24 h, the concentrations of STS were 10 / 20 / 30 / 40 / 50umol/L,hen the cell viability was measured by CCK-8 Kit.

### Caspase 3/7 was used to detect the apoptosis in the combination of STS and NaAsO2

The cells were divided into control group, NaAsO2 Group (15umol / L), STS group (20umol / L), NaAsO2 group and STS combined group, and CASPASE3 / 7 was detected by CASPASE3 / 7 Kit after 8 hours.

### Caspase 8 was used to detect the apoptosis of NaAsO2 after STS pretreatment

The cells were divided into Control Group, NaAsO2 group (10umol / L), STS was pretreated for 12h and NaAsO2 was added for 24 h,the concentrations of STS were 10 / 20 / 30 / 40 / 50umol/L, Caspase8 kit was used to detect the content of Caspase8 in cells.

### Statistical methods

SPSS 21. 0 Software was used to analyze the data, and univariate Anova was used to test the significance of the inter-group comparison. For two comparisons, either LSD-t method or Dunnett’s t 3 method was used when the homogeneity of the variance was satisfied. With a p < 0.05 was statistically significant.

## RESULTS

### The effect of STS alone on cell viability

Compared with the control group, different concentrations of STS had no significant effect on the cell viability (n = 6, P > 0.05), see figure 1.

**Fig 1:**
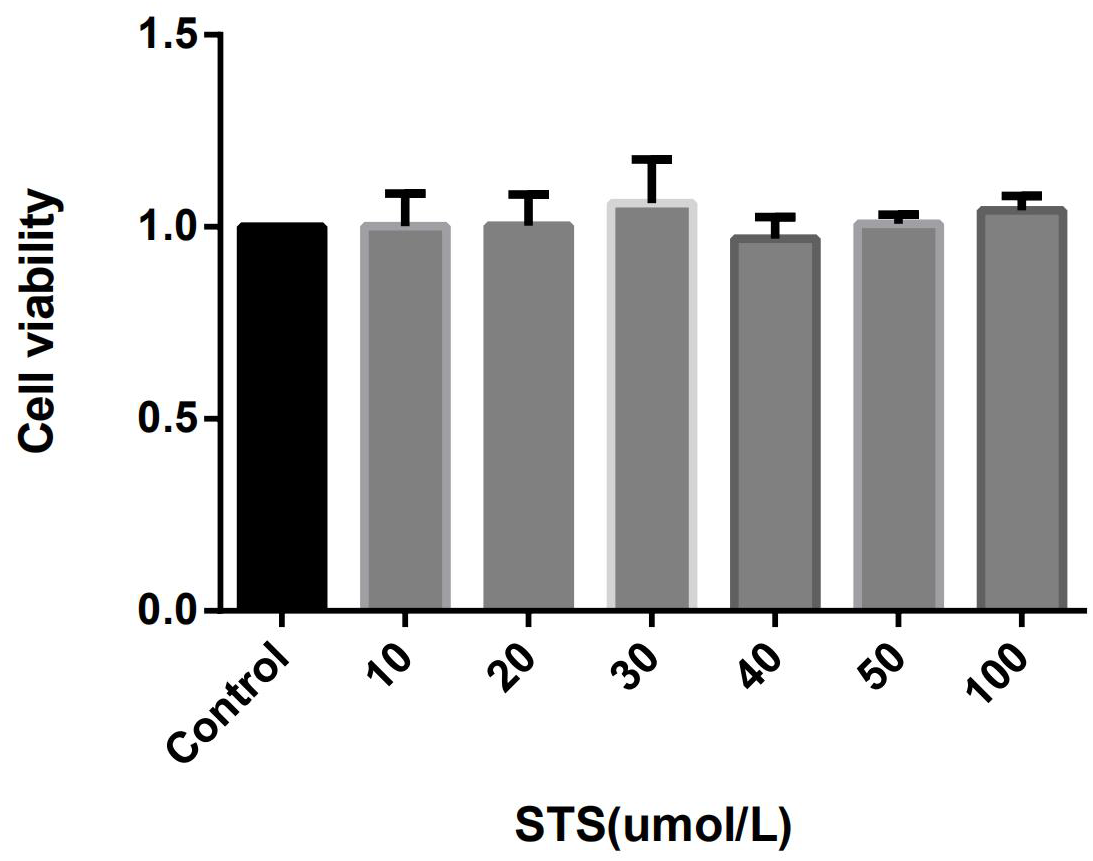
Effect of gradient concentration of STS on cell viability

### The effect of 6h after STS pretreatment and 12h after NaAsO2 treatment on cell viability

Compared with the control group, there was no significant change in the viability of cells treated with arsenic for 12 h after STS pretreatment for 6 h (n = 6, P > 0.05),the viability of cells treated with NaAsO2 alone was decreased(n = 6, P < 0.05), see figure 2.

**Fig 2:**
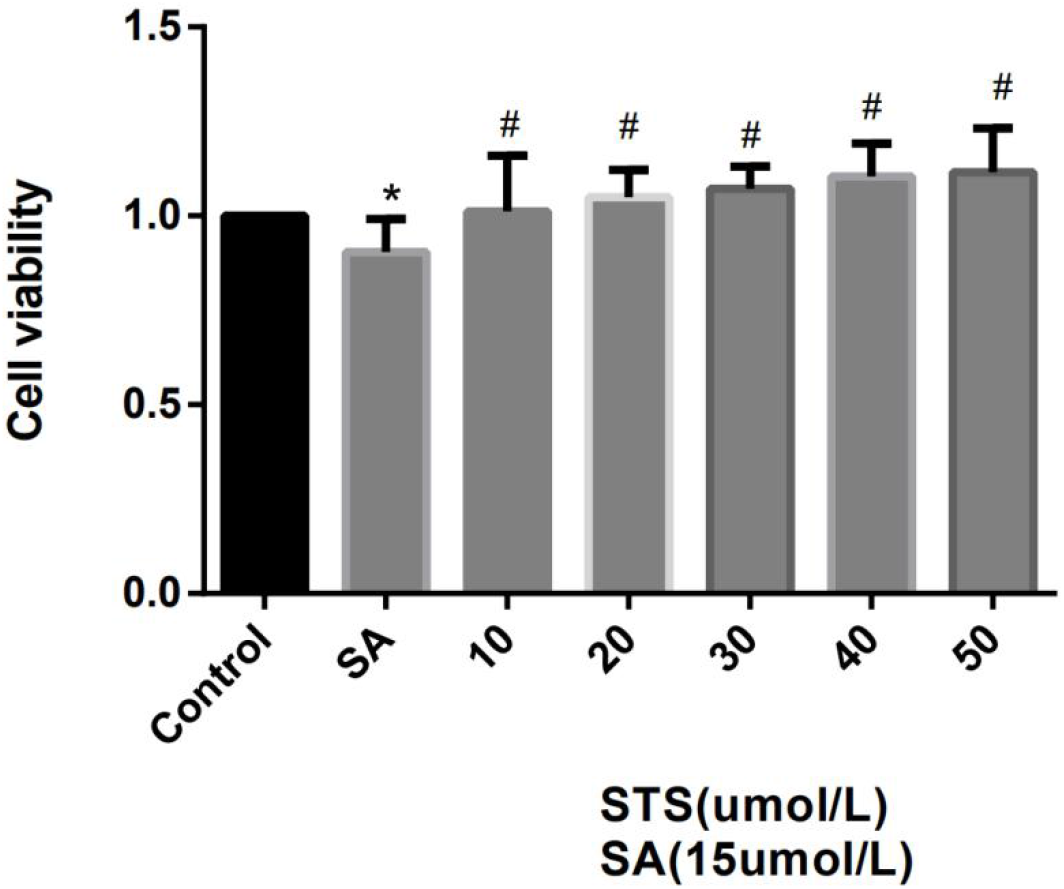
Changes of cell viability after STS pretreatment for 6 hours plus NaAsO2 12h n=6, Compared With Control Group, * p < 0.05 Compared with SA, # p < 0.05

### The effect of 12h after STS pretreatment and 24h after NaAsO2 treatment on cell viability

Compared with the control group, the cell viability was decreased after STS pretreatment for 12h and arsenic for 24h,the cell viability of NaAsO2 group was also decreased,the viability of NaAsO2 cells was lower than that of STS pretreated for 12h with NaAsO2 for 24h(n=6,p<0.05), see figure 3.

**Fig 3:**
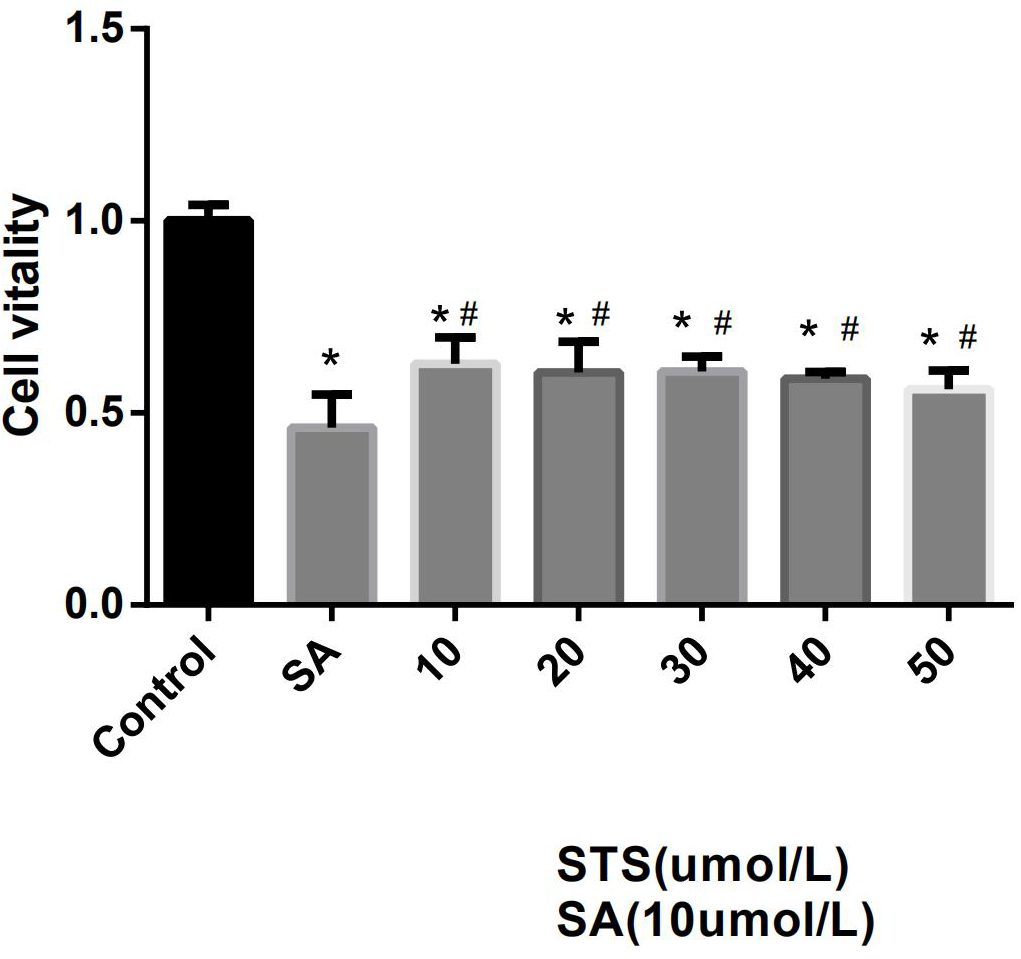
Changes of cell viability after STS pretreatment for 12 hours plus NaAsO2 24h Note: n=6, Compared With Control Group, * p < 0.05 Compared with SA, # p < 0.05

### Changes of Caspase3 / 7 in cells treated with combination of STS and NaAsO2

The content of Caspase 3/7 in the NaAsO2 group was higher than that in the STS and NaAsO2 group and the Control group and STS group after 12h,the differences were statistically significant(n=6,p<0.05), see figure 4.

**Fig 4:**
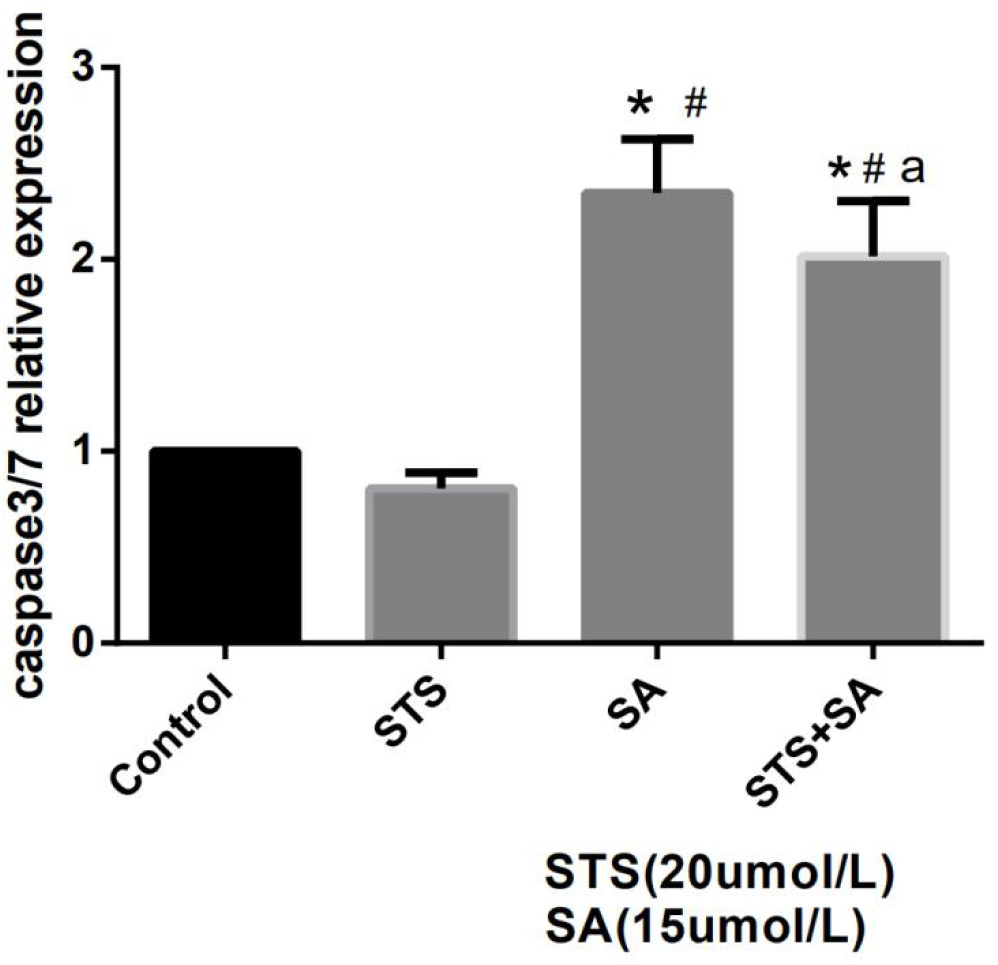
Relative Caspase 3 / 7 content after STS and NaAsO2 combined **treatment** n=6; Compared With Control Group, * p < 0.05 Compared with STS, # p < 0.05 Compared with SA, aP < 0.05

### Changes of Caspase8 in cells with 12h after STS pretreatment and 24h after NaAsO2 treatment

After STS pretreatment for 12h and NaAsO2 for 24 h, there was no significant difference in Caspase 8 content between NaAsO2 group and other groups (n=6,P>0.05), see figure 5

**Fig 5.**
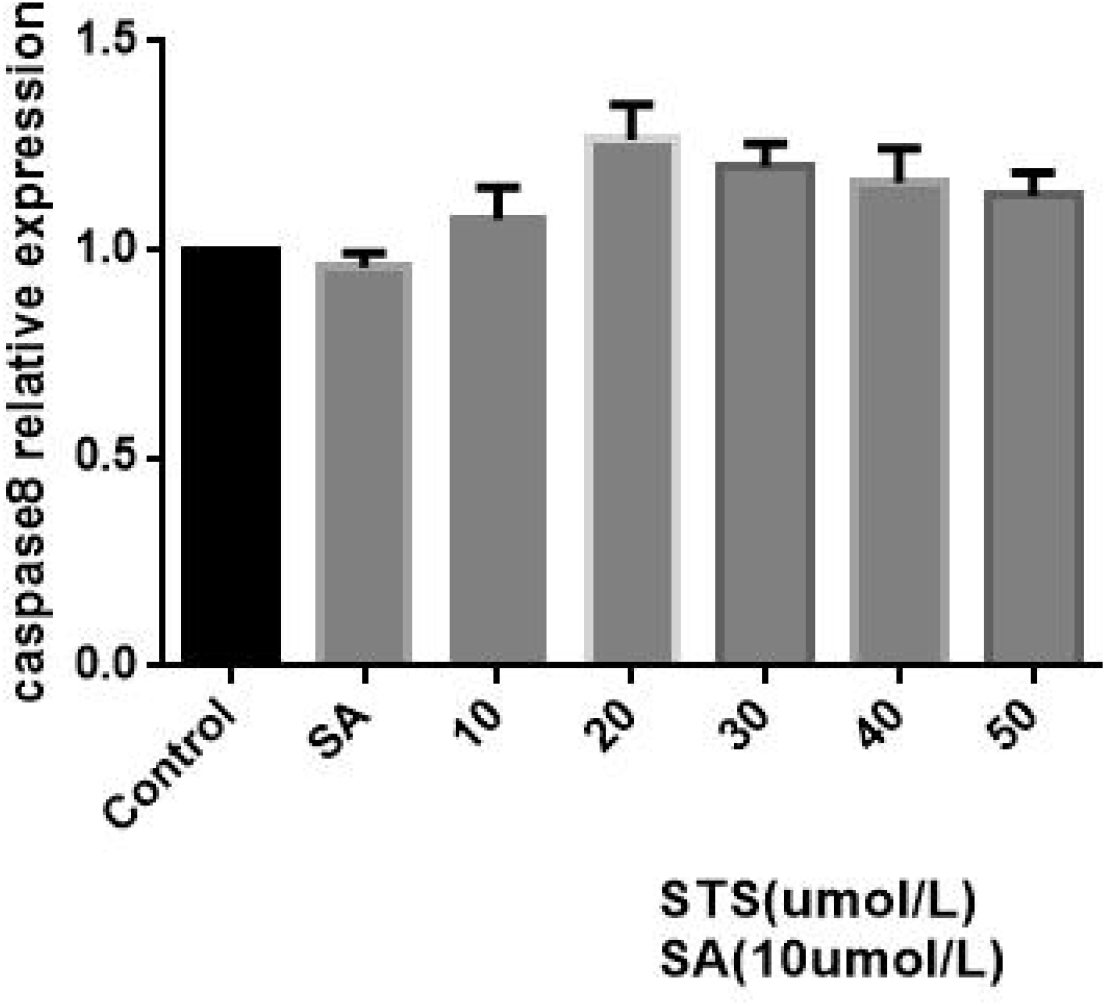
Caspase8 relative content after STS pretreatment plus NaAsO2.

## Discussion

Arsenic is an ancient toxic metal that is commonly found in the environment. NaAsO2 is the most toxic, affecting people in many countries around the world, including China.Arsenic has been shown to cause many cardiovascular diseases and to cause cardiomyocyte damage and cardiomyocyte apoptosis^5,6)^. Sodium tanshinone IIA sulfonate (STS) has been approved by China Food and Drug Administration (CFDA) for the treatment of cardiovascular disease.^7,8)^However, the combination of NaAsO2 poisoning and STS has not been reported. The effect of STS on myocardial injury induced by NaAsO2 was investigated at the cellular level.The results of CCK-8 Assay showed that A certain concentration of STS (0-50 and 100umol / l) had no cytotoxic effect on H9C2 cells. In addition, STS pretreatment can effectively reverse the decline in cell viability caused by NaAsO2 and reduce the apoptosis of H9c2 cardiomyocytes induced by sodium arsenite.

Caspase is a family of proteases that are closely related to apoptosis. They are mainly divided into initiating and executive types. Both are activated sequentially by foreign protein signals, leading to programmed cell death.The initial caspase-8 is a molecular switch for apoptosis and necrosis, and caspase-3 and -7 are related factors for performing apoptosis. Its overexpression has been shown to reduce myocardial contractility and cause heart disease in many pathological diseases.

Insufficiency.^.9.10,11,12)^Nagalakshmi Prasanna and other studies have confirmed that NaAsO2 administration can up-regulate caspase3 in rat heart tissue.^13^ and the results of this study are in good agreement with it.. In addition, STS can inhibit the expression of caspase3/7 in the NaAsO2-treated group in this study, which also suggests that STS may protect myocardial cells from the apoptosis induced by sodium arsenite to a certain extent.In this study, sodium tanshinone IIA sulfonate could inhibit the increase of caspase-3 expression level, but there was no significant change in the expression level of caspase-8. Previous studies have shown that Bax / Bcl-2, caspase-3, and caspase-9 play important roles in the mitochondrial apoptosis pathway, while caspase-8 and caspase-12 are death receptor signaling pathways and endoplasmic reticulum apoptosis pathways, respectively. Key factors in.^14)^Therefore, the anti-apoptotic effect of tanshinone IIA sodium sulfonate may be mediated through a mitochondrial-dependent apoptosis pathway.

In short, the use of sodium tanshinone IIA sulfonate can reverse the decline in cell viability and increase of apoptosis induced by sodium arsenite, which provides a feasible method for the inhibition of sodium arsenone IIA sulfonate on apoptosis The molecular explanation provides a new idea for the prevention and treatment of sodium arsenite-induced heart failure. Further research is needed to elucidate the exact mechanism of the protective effect of sodium tanshinone IIA sulfonate on myocardial apoptosis.

## ACKNOWLEDGMENTS

This work was supported in part by National Natural Science Foundation of China and Public Health Experimental Teaching Center of Shenyang Medical College.

